# Coupling Luciferase Activation and BRET Enables High-Contrast Red-Window Calcium Imaging

**DOI:** 10.64898/2026.07.26.740815

**Authors:** Rachna Sharma, Ying Xiong, Xiaodong Tian, Ponnusamy Pon Sathieshkumar, Hui-wang Ai

## Abstract

Genetically encoded bioluminescent calcium indicators enable excitation-free imaging of Ca^2+^ dynamics with low background, no photobleaching, and improved tissue penetration. However, many exhibit lower dynamic ranges than their fluorescent counterparts, motivating the development of new sensor configurations, particularly for imaging in the red optical window, where tissue absorption and scattering are reduced. Here, we developed HyBRIC and HyBRIC2, two hybrid indicators that integrate Ca^2+^-dependent luciferase activation and bioluminescence resonance energy transfer (BRET) within a single molecular architecture. When paired with pyCTZ luciferin, they exhibited an approximately 55-fold Ca^2+^-dependent signal increase in the red optical window *in vitro*. Compared with HyBRIC, HyBRIC2 showed improved brightness and responsiveness in mammalian cells. It robustly resolved induced Ca^2+^ dynamics in HEK293T and HeLa cells, with particularly strong red-window responses. Together, these findings expand the limited repertoire of responsive red-window bioluminescent calcium indicators and establish hybrid multiplicative sensing as a modular strategy for improving biosensor contrast.

---

Calcium ions (Ca^2+^) are ubiquitous second messengers that regulate diverse biological processes, including synaptic transmission, muscle contraction, gene expression, metabolism, and apoptosis.^1-4^ Imaging Ca^2+^ dynamics is therefore essential for understanding cellular signaling in living systems. Genetically encoded fluorescent calcium indicators have transformed Ca^2+^ imaging by enabling genetically targeted measurements in specific cells, organelles, and subcellular locations.^5-8^ However, fluorescence imaging requires external excitation light, which has limited tissue penetration and can introduce autofluorescence, phototoxicity, and photobleaching. External illumination may also interfere with photosensitive biological processes and optogenetic manipulations.^9,10^

Bioluminescent calcium indicators provide a complementary approach by generating photons through the luciferase-catalyzed oxidation of a luciferin.^11-13^ Because they do not require external illumination, bioluminescent indicators enable low-background imaging with minimal excitation-associated phototoxicity and photobleaching and are compatible with many light-sensitive biological processes.^11,14-16^ The development of compact and bright luciferases derived from *Oplophorus gracilirostris*, including NanoLuc and related engineered variants, has substantially expanded the performance and design space of bioluminescent calcium indicators.^17-19^ Nevertheless, many existing bioluminescent indicators exhibit more modest Ca^2+^-dependent signal changes than leading fluorescent indicators. Developing new molecular configurations and signal-transduction mechanisms that increase their dynamic range therefore remains an important objective.

Bioluminescent calcium indicators can be broadly classified according to the primary optical mechanism by which Ca^2+^ binding generates a measurable response. In intensiometric indicators, Ca^2+^-dependent conformational changes modulate luciferase activity, thereby altering total photon output.^8,18,20^ This strategy has been implemented by inserting Ca^2+^-responsive domains into luciferases or by coupling Ca^2+^ binding to luciferase complementation or reconstitution. Representative examples include GeNL(Ca^2+^),^21^ CaMBI,^18^ BRIC,^19^ (**Fig. S1A**), CaBLAM,^22^ and related indicator families. Some of these indicators incorporate a long-wavelength-emitting fluorescent protein as a constitutive bioluminescence resonance energy transfer (BRET) acceptor to red-shift the emission;^18,19^ however, the Ca^2+^-dependent signal change arises primarily from modulation of luciferase activity rather than from changes in BRET efficiency (**Fig. S1A**). In the second major class, a luciferase donor is paired with a fluorescent-protein-derived acceptor, and Ca^2+^ binding alters BRET efficiency. CalfluxVTN is a representative example in which a Ca^2+^-binding troponin C (TnC) domain changes the relative geometry of the donor and acceptor, thereby modulating BRET efficiency and producing a Ca^2+^-dependent ratiometric emission response (**Fig. S1B**).^14,23^ Several other studies have expanded these two general strategies by combining luciferases with genetically encoded or synthetic fluorescent Ca^2^□ sensors, yielding indicators such as LUCI-GECO1,^24^ GLICO,^20^ and H-Luc-MaPCa.^25^ Together, these architectures illustrate the diverse mechanisms through which Ca^2+^ binding can regulate bioluminescence output.

Emission wavelength is another critical consideration for biological imaging. Blue and green photons are strongly affected by tissue absorption and scattering, whereas longer-wavelength photons generally exhibit improved transmission through biological tissues.^12,19,26^ Red-shifted emission above 600 nm is therefore particularly desirable for longitudinal and *in vivo* bioluminescence imaging. Despite recent progress in bioluminescent calcium indicator development, relatively few indicators combine robust Ca^2+^-dependent signal changes with substantial emission in the red optical window (**Table S1**). New sensor architectures that selectively amplify Ca^2+^-dependent emission at these wavelengths could therefore improve the sensitivity and applicability of bioluminescent calcium imaging.

We reasoned that Ca^2+^-dependent luciferase activation and Ca^2+^-dependent BRET could be integrated within a single molecular architecture to enhance red-window contrast (**Fig. S1C**). Coupling these two Ca^2+^-responsive processes could allow their optical effects to reinforce one another, producing a multiplicative increase in red-window emission. Here, we report the development of two hybrid bioluminescent calcium indicators, HyBRIC and HyBRIC2, by combining a CaM-M13 regulated luciferase insertion module with a TnC-regulated BRET module. By optimizing the individual modules and their molecular coupling and comparing luciferin substrates, we identified pyCTZ as the luciferin that best balanced photon output and energy transfer while minimizing donor emission leakage into the red optical window. The resulting indicators exhibited greater than 55-fold Ca^2+^-dependent signal changes in the red optical window *in vitro*. Although the first-generation indicator showed a large *in vitro* response, its response was more limited in mammalian cells. Further engineering of HyBRIC into HyBRIC2 resulted in increased brightness and improved responses in living cells. HyBRIC2 resolved induced Ca^2+^ dynamics in HEK293T and HeLa cells, with particularly strong responses in the red optical window. Together, these results demonstrate that coupling luciferase activation and BRET provides a modular hybrid strategy for enhancing the contrast of bioluminescent biosensors.

## EXPERIMENTAL SECTION

### General information

DNA oligonucleotides used in this study were synthesized by Integrated DNA Technologies (IDT) or Eurofins. Restriction endonucleases, high-fidelity DNA polymerase, and Taq DNA polymerase were obtained from Thermo Fisher Scientific or New England Biolabs. PCR products and restriction-digested DNA fragments were separated by gel electrophoresis and purified using Syd Laboratories Gel Extraction columns. Plasmid DNA was isolated from pellets of 5 mL overnight LB cultures by alkaline lysis using Syd Laboratories Miniprep columns. DNA sequencing was performed by Eurofins. The luciferins pyCTZ, DTZ, and 8pyDTZ were prepared according to previously reported synthetic procedures.^12,27^

### Plasmids and library construction

To optimize the Ca^2+^-dependent BRET response (Pathway 1; **Fig. 1A**), a gene fragment encoding CalfluxVTN^14^ was synthesized and cloned into a pBAD expression vector between the XhoI and HindIII restriction sites. CaRET0.2 was subsequently generated by replacing the fluorescent-protein acceptor and luciferase donor with a C-terminally truncated mScarlet-I^28^ and LumiLuc,^12^ respectively, using Gibson assembly. Linker-mutagenesis libraries were then generated using oligonucleotides containing degenerate codons at the corresponding residue positions. NNK and NDT codons were used, where N represents A, C, G, or T; K represents G or T; and D represents A, G, or T. To optimize Ca^2+^-dependent luciferase activation (Pathway 2; **Fig. 1B**), a CaM-M13 sensing domain was inserted into LumiScarlet, a genetic fusion of mScarlet-I and LumiLuc.^12^ The CaM-M13 coding sequence was amplified from pcDNA3.1-Orange_CaMBI_110 (Addgene plasmid #124094)^18^ and inserted at residue 66 or 133 of LumiLuc in a pBAD construct derived from Addgene plasmid #126623 using multi-fragment Gibson assembly. Insertion at residue 66 using linker sequences derived from Orange CaMBI or GeNL(Ca^2+^) yielded L-BRIC66a and L-BRIC66b, respectively. Insertion at residue 133 using the Orange CaMBI-derived linker configuration yielded L-BRIC133. The first-generation hybrid plasmid, pBAD-HyBRIC, was constructed by combining the optimized CaRET and L-BRIC133 components within a single coding sequence using multifragment Gibson assembly. pBAD-HyBRIC was subsequently used as the template for iterative rounds of error-prone PCR mutagenesis to generate libraries from which HyBRIC2 was isolated. For mammalian expression, DNA fragments encoding HyBRIC and HyBRIC2 were amplified from the corresponding pBAD plasmids and inserted into XhoI/HindIII-digested pcDNA3 by Gibson assembly.

**Figure 1.**
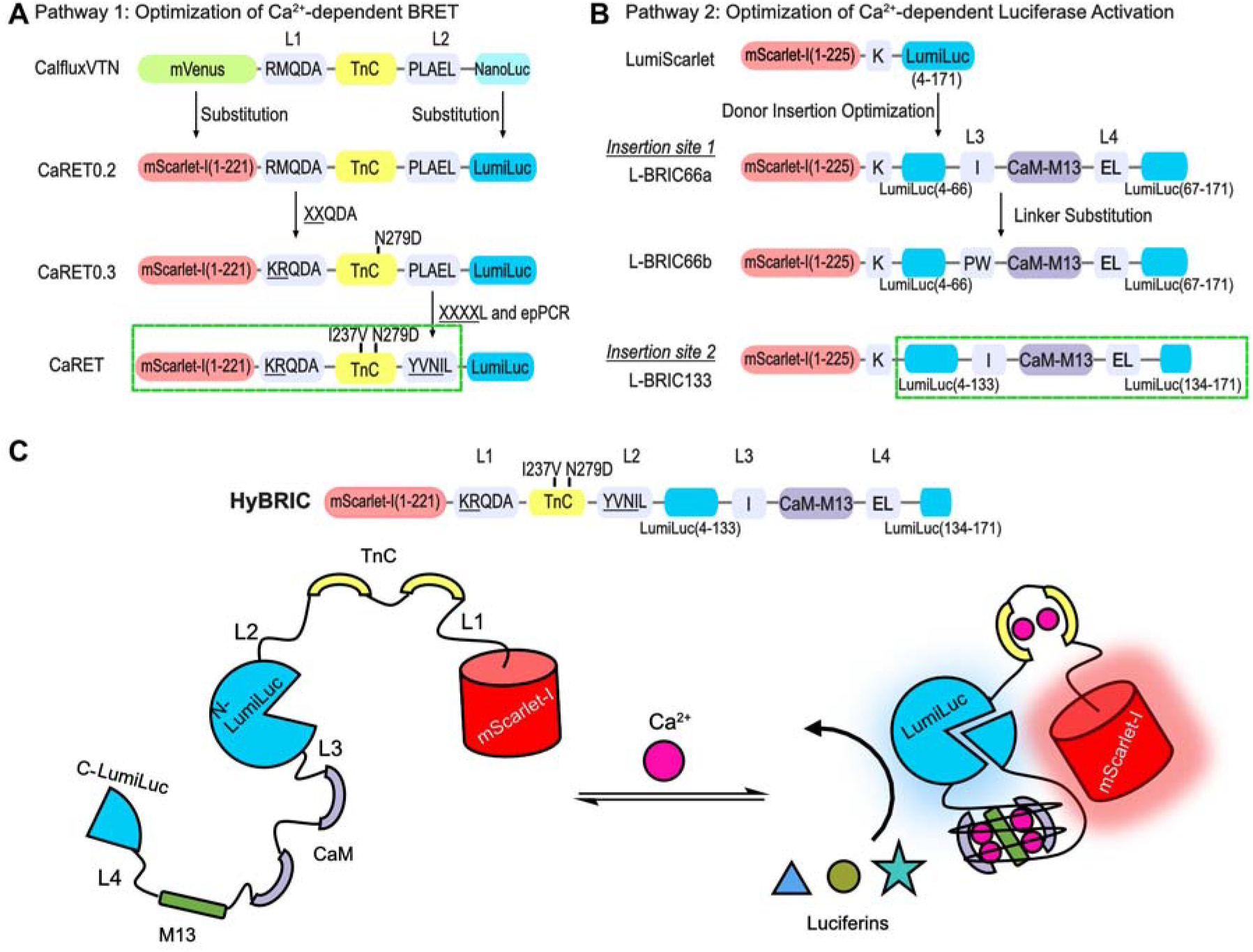
Modular design and dual-pathway optimization of the hybrid bioluminescent calcium indicator HyBRIC. (**A**) Engineering workflow for optimizing Ca^2+^-dependent BRET modulation (Pathway 1). Sequential donor and acceptor replacement, linker optimization, and random mutagenesis yielded CaRET, a ratiometric bioluminescent Ca^2+^ indicator exhibiting a 12-fold Ca^2+^-dependent change in the BRET ratio. (**B**) Engineering workflow for optimizing Ca^2+^-dependent luciferase activation (Pathway 2). The CaM-M13 sensing domain was inserted at different positions within LumiLuc, followed by linker optimization. Among the tested variants, L-BRIC133 exhibited the largest response, with a 4.9-fold Ca^2+^-dependent increase in bioluminescence intensity. (**C**) Domain organization and proposed sensing mechanism of HyBRIC. The optimized CaRET and L-BRIC133 modules, outlined by green boxes in panels A and B, respectively, were combined within a single molecular scaffold. Ca^2+^ binding promotes both split-LumiLuc reconstitution and TnC-mediated enhancement of BRET to mScarlet-I, resulting in a coupled increase in red-window bioluminescence.

### Phenotypic screening and *in vitro* characterization

For library screening, *Escherichia coli* DH10B cells transformed with the corresponding plasmid libraries were plated on LB agar containing ampicillin (100 μg/mL) and L-arabinose (0.1%, w/v) and incubated overnight at 37 °C. Before imaging, the plates were equilibrated at room temperature for 6 h. Bioluminescence imaging was performed in a UVP BioSpectrum dark box equipped with a QSI 628 cooled CCD camera (Quantum Scientific Imaging). Approximately 500 μL of 1 mM CaCl_2_ was applied uniformly to each plate, followed by incubation in the dark box for 5 min. Approximately 500 μL of 50 μM pyCTZ was then applied. Bioluminescence images were acquired through a 600 nm long-pass filter and analyzed using Fiji/ImageJ. The 192 brightest colonies were selected and transferred to deep-well 96-well plates containing 500 μL of LB medium supplemented with ampicillin (100 μg/mL) and L-arabinose (0.1%, w/v) per well. After overnight growth at 37 °C with shaking at 250 rpm, the cultures were centrifuged, the supernatants were removed, and the cell pellets were lysed with 500 μL of B-PER reagent (Thermo Fisher Scientific). For secondary screening, 5 μL of each lysate was added to either 185 μL of Ca^2+^-free buffer (30 mM MOPS, 100 mM KCl, and 10 mM EGTA, pH 7.2) or 185 μL of another buffer with 39 μM free Ca^2+^ (30 mM MOPS, 100 mM KCl, and 10 mM CaEGTA, pH 7.2). 10 μL of a 400 μM luciferin stock solution was injected into each well using the reagent injector of a CLARIOstar Plus microplate reader (BMG Labtech). Bioluminescence emission spectra were recorded from 400 to 800 nm in 5 nm increments using the red-wavelength-extended photomultiplier tube. The same secondary-screening workflow was used during the evolution of HyBRIC2, except that variants were compared at 65 nM and 1.35 μM free Ca^2+^.^26^ Bacterial lysates prepared from three independently cultured colonies were used for spectral and Ca^2+^-affinity characterization of HyBRIC and HyBRIC2 by following published procedures.^19,26^ Data were analyzed using GraphPad Prism 10. Apparent dissociation constants and Hill coefficients were determined by fitting the data to a four-parameter Hill equation.

### Characterization of HyBRIC and HyBRIC2 in mammalian cells

HEK293T cells (ATCC, Cat. No. CRL-3216) and HeLa cells (ATCC, Cat. No. CCL-2) were transfected with 3 μg of pcDNA3-HyBRIC or pcDNA3-HyBRIC2 per 35 mm dish. Following overnight incubation in high-glucose DMEM containing 4.5 g/L glucose at 37 °C in a humidified 5% CO□ incubator, the cells were rinsed twice with DPBS containing 0.9 mM Ca^2+^ and incubated in the same buffer for 15 min before imaging. DPBS containing 0.9 mM Ca^2+^ was used as the imaging buffer throughout the experiments. Images were acquired using an inverted Leica DMi8 microscope equipped with a Photometrics Prime 95B scientific CMOS camera. The microscope was housed in a dark enclosure and controlled using Leica LAS X software, version 3.5.7. Imaging was performed using a 40× oil-immersion objective with a numerical aperture of 1.2 and without a filter cube. Camera settings were as follows: 2 × 2 binning, 10 s exposure time, 0 s interval, −20 °C sensor temperature, 12-bit acquisition, and high-sensitivity mode. No emission filter condition was used to acquire the total bioluminescence, while an RFP imaging filter cube (with a 610/75 nm bandpass filter) was used to collect red window bioluminescence. Images were processed and analyzed using ImageJ/Fiji according to established protocols.^19^ To initiate bioluminescence imaging, the imaging buffer was replaced with fresh DPBS containing 50 μM pyCTZ. After a stable baseline was recorded for 4-5 frames, acetylcholine (Thermo Scientific, Cat. # AC159170050) was added to a final concentration of 10 μM to stimulate intracellular Ca^2+^ mobilization. For measurements of Ca^2+^ oscillations in HeLa cells, histamine (Chem-Impex, Cat. # 21888) dissolved in DPBS was added during time-lapse imaging to a final concentration of 20 μM after the addition of 50 μM pyCTZ.

## RESULTS

### Stepwise Development of HyBRIC

To develop bioluminescent Ca^2+^ indicators with large responses in the red optical window, we explored a hybrid strategy that integrates Ca^2+^-dependent modulation of luciferase activity and BRET within a single construct. In this design (**Fig. S1C**), two Ca^2+^-responsive modules perform distinct functions: a CaM-M13 module regulates luciferase catalytic activation, whereas a TnC-based module alters donor-acceptor geometry and BRET efficiency. The two processes are combined to maximize Ca^2+^-dependent optical changes in the optical window: increased photon production through catalytic reconstitution and enhanced red-window emission through BRET-mediated spectral redistribution. We therefore used a bifurcated engineering strategy in which the ratiometric BRET pathway and the intensiometric catalytic pathway were optimized separately before being integrated into a unified scaffold. We utilized the pyCTZ luciferin for sensor screening as it was shown to be one of the brightest LumiLuc substrates.^12^

We first optimized the BRET-based Ca^2+^ response (**Fig. 1A**). Inspired by CalfluxVTN^14^ and our previous use of a C-terminally truncated mScarlet-I as the BRET acceptor in BRIC,^19^ we replaced the fluorescent protein in CalfluxVTN with the same truncated mScarlet-I variant comprising residues 1-221 and replaced the luciferase donor with LumiLuc. LumiLuc was selected because of its ability to efficiently utilize a range of luciferins.^12^ The resulting construct, CaRET0.2, exhibited an approximately 3.6-fold Ca^2+^-dependent change in the BRET ratio (R/R_0_; **Fig. S2**). We next optimized linker 1 (L1; **Fig. 1A**) by introducing two randomized codons. Screening the resulting library in bacterial lysates identified CaRET0.3, which exhibited an approximately 5.5-fold Ca^2+^-dependent change in the BRET ratio (**Fig. S2**). In addition to the L1 mutations, CaRET0.3 contained a serendipitous N279D mutation in the TnC domain, likely gained during PCR amplification. We then introduced random mutations into linker 2 (L2), followed by error-prone PCR-based mutagenesis. This iterative optimization yielded the CaRET variant with an approximately 12-fold Ca^2+^-dependent ratiometric change (R/R_0_) *in vitro*.

In parallel, we optimized Ca^2+^-dependent catalytic activation by inserting a CaM-M13 sensing domain into LumiScarlet,^12^ a genetic fusion of mScarlet-I and LumiLuc (**Fig. 1B**). Two insertion sites, at residues 66 and 133 of LumiLuc, were selected based on architectures previously used in Orange CaMBI and GeNL(Ca^2+^).^14,21^ Using the CaM-M13 module and linker composition adopted from Orange CaMBI, insertion at residues 66 and 133 generated L-BRIC66a and L-BRIC133, respectively. L-BRIC66a exhibited an approximately 1.4-fold Ca^2+^-dependent increase in bioluminescence (BL/BL_0_), whereas L-BRIC133 showed a substantially larger response of approximately 4.9-fold under Ca^2+^-saturating conditions. We also tested the linker composition used in GeNL(Ca^2+^) at residue 66, generating L-BRIC66b, which exhibited an approximately 3.4-fold response (**Fig. S2**). We therefore selected L-BRIC133 as the optimized catalytic module and combined it with the optimized BRET module, CaRET, within a single construct to generate the first-generation hybrid indicator, HyBRIC (**Figs. 1C and S3**).

### Substrate Profiling and *In Vitro* Characterization for HyBRIC

Because the hybrid architecture relies on both luciferase donor emission and BRET-mediated mScarlet-I acceptor emission, substrate selection required balancing donor brightness, spectral overlap with the acceptor, and minimal donor leakage into the red window. Previous studies have shown that LumiLuc has broad substrate compatibility and can efficiently use a group of luciferins with distinct spectral properties.^12,27^ We therefore compared the emission profiles of HyBRIC paired with pyCTZ, DTZ, or 8pyDTZ under Ca^2+^-free and Ca^2+^-saturated conditions (**Fig. 2**).

**Figure 2.**
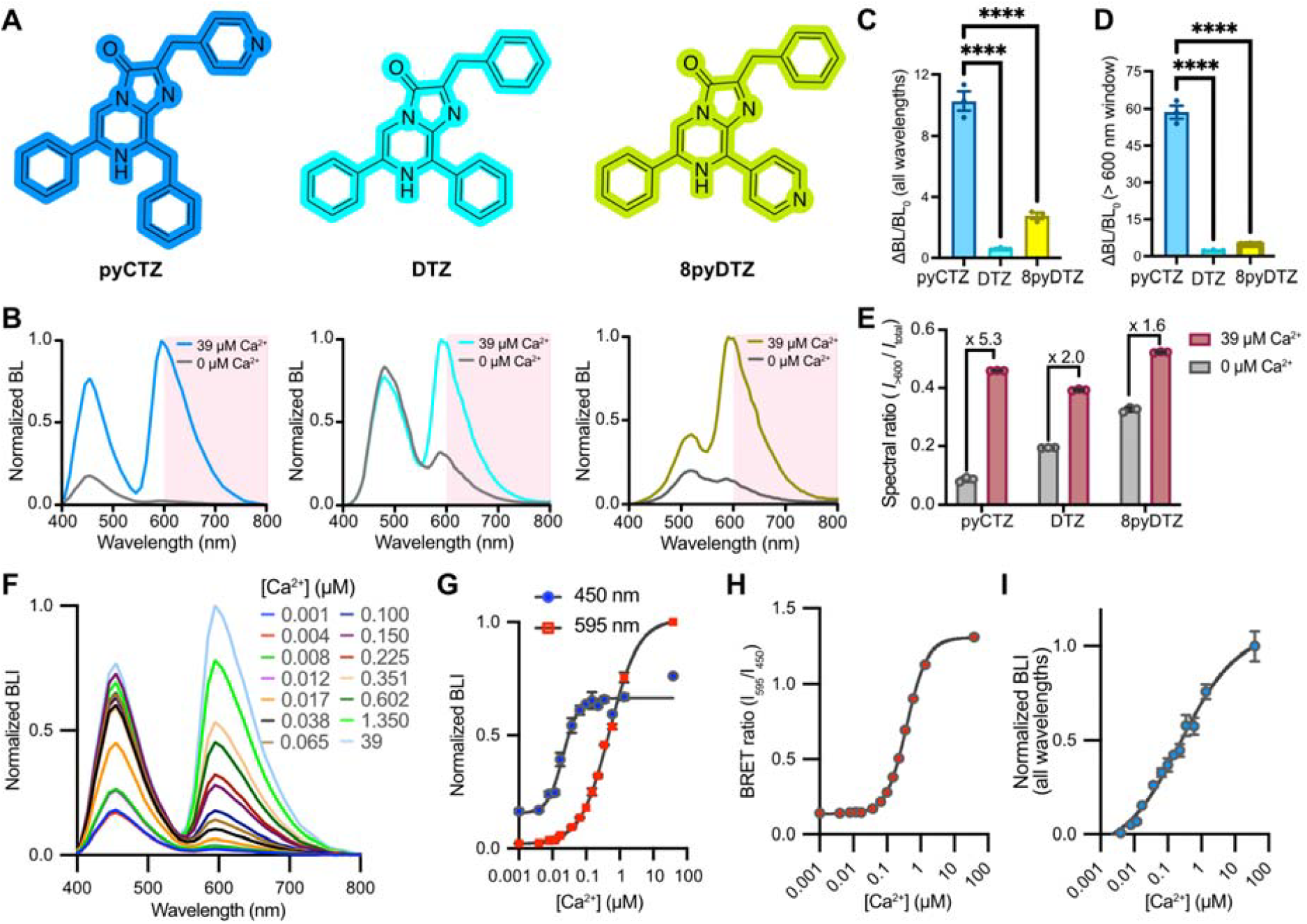
Substrate profiling and *in vitro* characterization of HyBRIC. (**A**) Chemical structures of pyCTZ, DTZ, and 8pyDTZ. (**B**) Normalized bioluminescence emission spectra of HyBRIC with each luciferin under Ca^2+^-free (0 μM) and Ca^2+^-saturated (39 μM) conditions. The red optical window (λ > 600 nm) is shaded. (**C**) Relative change (ΔBL/BL□) in total bioluminescence integrated from 400 to 800 nm. (D) Relative change (ΔBL/BL□) in red-window bioluminescence integrated from 600 to 800 nm. (E) Spectral ratios, calculated as the integrated emission above 600 nm divided by the total integrated emission from 400 to 800 nm (*I*□_>600 nm_□/*I*_total_), under Ca^2+^-free and Ca^2+^-saturated conditions. The corresponding Ca^2+^-dependent changes in the spectral ratio are indicated above the brackets. (**F**) Normalized bioluminescence emission spectra of HyBRIC in bacterial lysates over a range of free Ca^2+^ concentrations. (**G**) Ca^2^□ titration curves based on normalized donor emission at 450 nm and acceptor emission at 595 nm, revealing stepwise activation of the two Ca^2+^ -responsive modules. Four-parameter Hill fitting yielded apparent *K*_d_ values of 21 nM and 490 nM and Hill coefficients of 1.9 and 1.0 for the 450- and 595-nm responses, respectively. (**H**) Ca^2+^ titration curve based on the BRET ratio (*I*_595_/*I*_450_), yielding an apparent *K*_d_ of 383 nM and a Hill coefficient of 1.4. (**I**) Ca^2+^ titration curve based on total bioluminescence integrated from 400 to 800 nm, yielding an apparent *K*_d_ of 238 nM and a Hill coefficient of 0.5. For panels C-E and G-I, data are presented as mean ± SEM from *n* = 3 replicates. Statistical significance was determined by ordinary one-way ANOVA followed by multiple-comparisons testing; ****P < 0.0001.

HyBRIC exhibited distinct spectral profiles depending on the substrate used. As expected, DTZ and 8pyDTZ produced more red-shifted emission than pyCTZ. Although this shift increased spectral overlap with mScarlet-I and enhanced BRET efficiency, it also increased donor-emission bleedthrough into the red optical window (**Fig. 2B-E**). Among the substrates tested, pyCTZ provided the greatest Ca^2+^-dependent contrast because it produced minimal basal red-window emission and limited donor bleedthrough. Under Ca^2+^-saturating conditions, HyBRIC paired with pyCTZ exhibited an approximately 11-fold increase in total integrated bioluminescence and an approximately 55-fold increase in red-window bioluminescence (ΔBL/BL_0_; **Fig. 2C,D**).

We next characterized the Ca^2+^ response of HyBRIC using *in vitro* titrations spanning 0-39 μM free Ca^2+^. Ca^2+^ produced concentration-dependent increases in both donor emission at approximately 450 nm and acceptor emission at approximately 595 nm, but the two channels responded over distinct Ca^2+^ concentration ranges. The donor-associated response had an apparent *K*_d_ of approximately 21 nM, whereas the acceptor-associated response had an apparent *K*_d_ of approximately 490 nM (**Fig. 2F,G**). In addition, the Ca^2^□-dependent BRET-ratio response had an apparent *K*_d_ of approximately 383 nM, whereas the response calculated from total bioluminescence integrated from 400 to 800 nm had an apparent *K*_d_ of approximately 238 nM (**Fig. 2H,I**).

This approximately 23-fold difference in apparent Ca^2+^ sensitivity at 450 nm and 595 nm supports stepwise activation of the two sensing modules as the Ca^2+^ concentration increases. At lower Ca^2+^ concentrations, the higher-affinity CaM-M13 module promotes split-luciferase reconstitution and increases photon production. At higher Ca^2+^ concentrations, the TnC-mediated conformational transition enhances BRET and redistributes the emitted light toward the red optical window. There is a narrow, overlapping intermediate concentration range, where both mechanisms are sensitive to Ca^2+^ concentration changes (**Fig. 2G**).

### Live-Cell Characterization and Engineering of HyBRIC2

To evaluate sensor performance in living cells, HyBRIC was transiently expressed in HEK293T cells. Following substrate addition, cells were stimulated with 10 μM acetylcholine (ACh), which activates G protein-coupled receptors and induces an increase in cytosolic Ca^2+^. Changes in both total bioluminescence and emission within the red window were monitored. Although HyBRIC remained responsive in mammalian cells, its cellular performance was substantially lower than expected from its *in vitro* characterization. ACh stimulation produced an approximately 41% increase in total bioluminescence and an approximately 72% increase in red-window emission (**Fig. 3A,B**). In our prior bacterial expression experiments, a substantial fraction of HyBRIC was noticed in the insoluble inclusion body fraction, which displayed the red color of mScarlet-I. These observations suggested that the complex multidomain construct may be prone to inefficient protein folding. In addition, the high Ca^2+^ affinity of the CaM-M13 regulated catalytic module may have caused partial activation under resting cellular conditions. The donor-associated apparent *K*_d_ of approximately 21 nM may have placed a substantial fraction of HyBRIC in a preactivated state before stimulation, thereby reducing the available inactive sensor population and compressing its response to stimulated Ca^2+^ transients.

**Figure 3.**
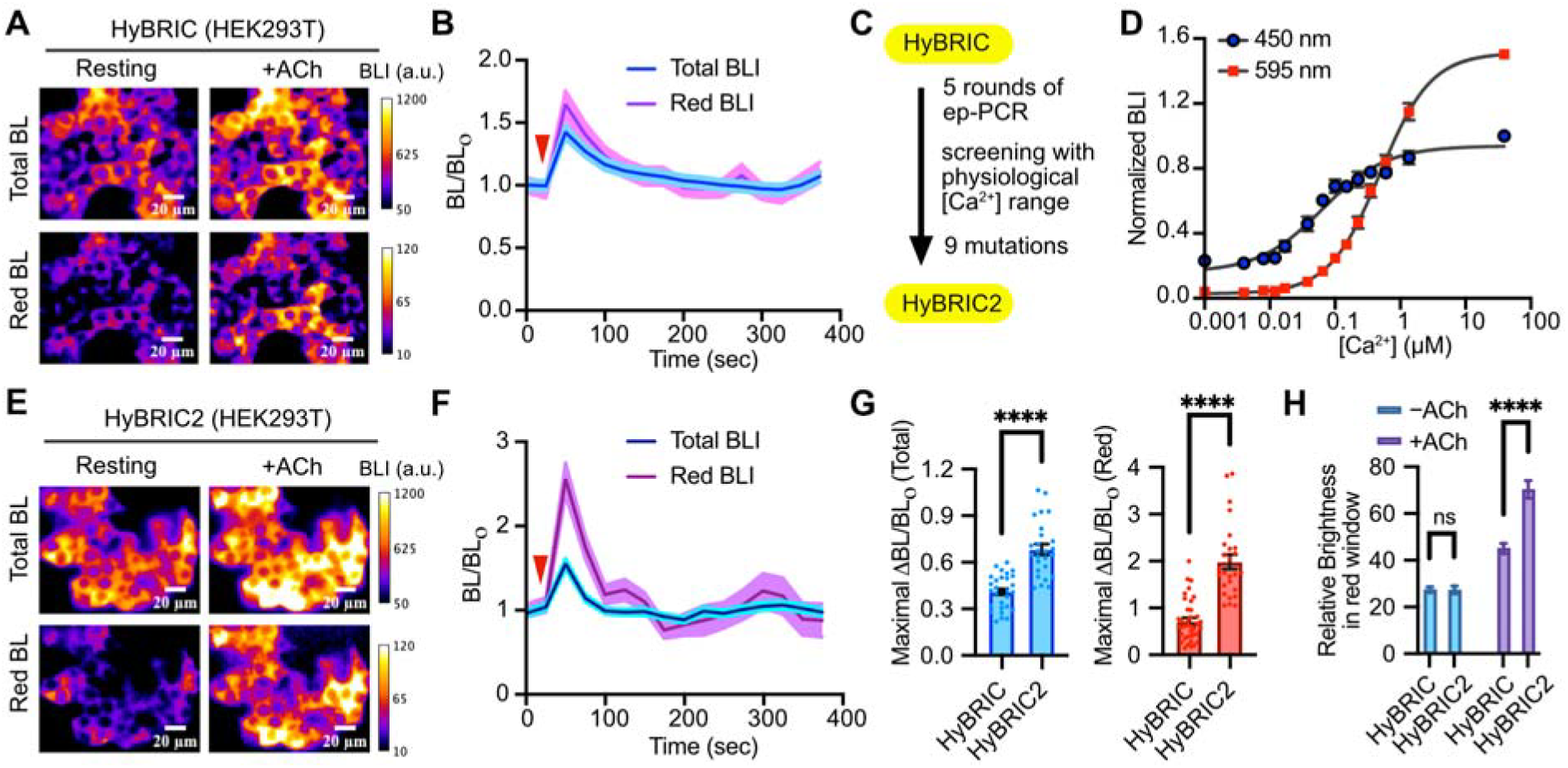
Live-cell characterization and directed evolution of HyBRIC into HyBRIC2. (**A**) Representative pseudocolored images of total and red-window bioluminescence intensity (BLI) from HEK293T cells transiently expressing HyBRIC before and after stimulation with 10 μM acetylcholine (ACh). Scale bars, 20 μm. (**B)** Time courses of total and red-window bioluminescence from HyBRIC-expressing cells, normalized to the pre-stimulation baseline (BL/BL_0_). The red triangle indicates the time of ACh addition. (**C**) Directed-evolution workflow used to generate HyBRIC2 from HyBRIC. (**D**) *In vitro* Ca^2+^ titration curves of HyBRIC2 in bacterial lysates, based on normalized donor emission at 450 nm and acceptor emission at 595 nm. Four-parameter Hill fitting yielded apparent *K*_d_ values of 64 and 489 nM and Hill coefficients of 0.9 and 1.1 for the 450- and 595-nm responses, respectively. (**E**) Representative pseudocolored images of total and red-window BLI from HEK293T cells transiently expressing HyBRIC2 before and after stimulation with 10 μM ACh. Scale bars, 20 μm. (**F**) Time courses of total and red-window bioluminescence from HyBRIC2-expressing cells, normalized to the pre-stimulation baseline. The red triangle indicates the time of ACh addition. (**G**) Comparison of the maximal fractional responses (ΔBL/BL_0_) of HyBRIC and HyBRIC2 based on total bioluminescence (left) and red-window bioluminescence (right). (**H**) Comparison of the relative red-window brightness of HyBRIC and HyBRIC2 under resting (−ACh) and stimulated (+ACh) conditions. Data in panel D are presented as mean ± SEM from *n* = 3 replicates. In panels B and F, solid lines and shaded regions represent mean ± SEM. Data in panels G and H are presented as mean ± SEM from *n* = 37 cells for HyBRIC and *n* = 39 cells for HyBRIC2; individual points in panel H represent single cells. Statistical significance was determined using unpaired two-tailed *t* tests; ns, not significant; ****P < 0.0001.

To improve live-cell performance, we subjected HyBRIC to iterative rounds of error-prone (EP) PCR-based directed evolution (**Fig. 3C**). Whereas the initial engineering of HyBRIC used 0 and 39 μM free Ca^2+^, subsequent libraries were screened at 65 nM and 1.35 μM free Ca^2+^ to better distinguish variants across a physiologically relevant Ca^2+^ range.^26^ This process yielded HyBRIC2 (**Fig. S3)**. Nine mutations were obtained, which are scattered through the mScarlet-I (2 mutations), TnC (3 mutations), N-terminal LumiLuc (1 mutation), and CaM-M13 (3 mutations) fragments (**Fig. S4**). Relative to HyBRIC, HyBRIC2 showed a shift in the donor-associated apparent *K*_d_ at 450 nm from approximately 21 to 64 nM while largely retaining the other photophysical and response properties (**Figs. 3D and S4**). This recalibration substantially improved live-cell performance. Following the same ACh stimulation, HyBRIC2 exhibited an approximately 68% increase in total bioluminescence and an approximately 200% increase in red-window emission (ΔBL/BL_0_; **Fig. 3E-G**). In addition, in the post-stimulation condition, HyBRIC2 was notably brighter than HyBRIC2 (**Fig. 3H**). Compared with the first-generation HyBRIC construct, HyBRIC2 therefore provided higher brightness and greater contrast, particularly in the red window, for imaging Ca^2+^ transients in mammalian cells.

We next examined whether HyBRIC2 could resolve endogenous Ca^2+^ dynamics in another mammalian cell type. HeLa cells expressing HyBRIC2 were stimulated with 20 μM histamine, a well-established trigger of intracellular Ca^2+^ oscillations. HyBRIC2 resolved successive oscillatory Ca^2+^ transients in both total integrated bioluminescence and emission within the red optical window (**Fig. 4**). The red-window signal consistently exhibited greater contrast than total emission, demonstrating the advantage of coupling catalytic activation with BRET-mediated spectral redistribution. Peak red-window responses reached up to an average of 400% (ΔBL/BL_0_) while the maximal response was observed to nearly 10-fold, enabling clearer visualization of Ca^2+^ oscillations than measurements based on total bioluminescence alone.

**Figure 4.**
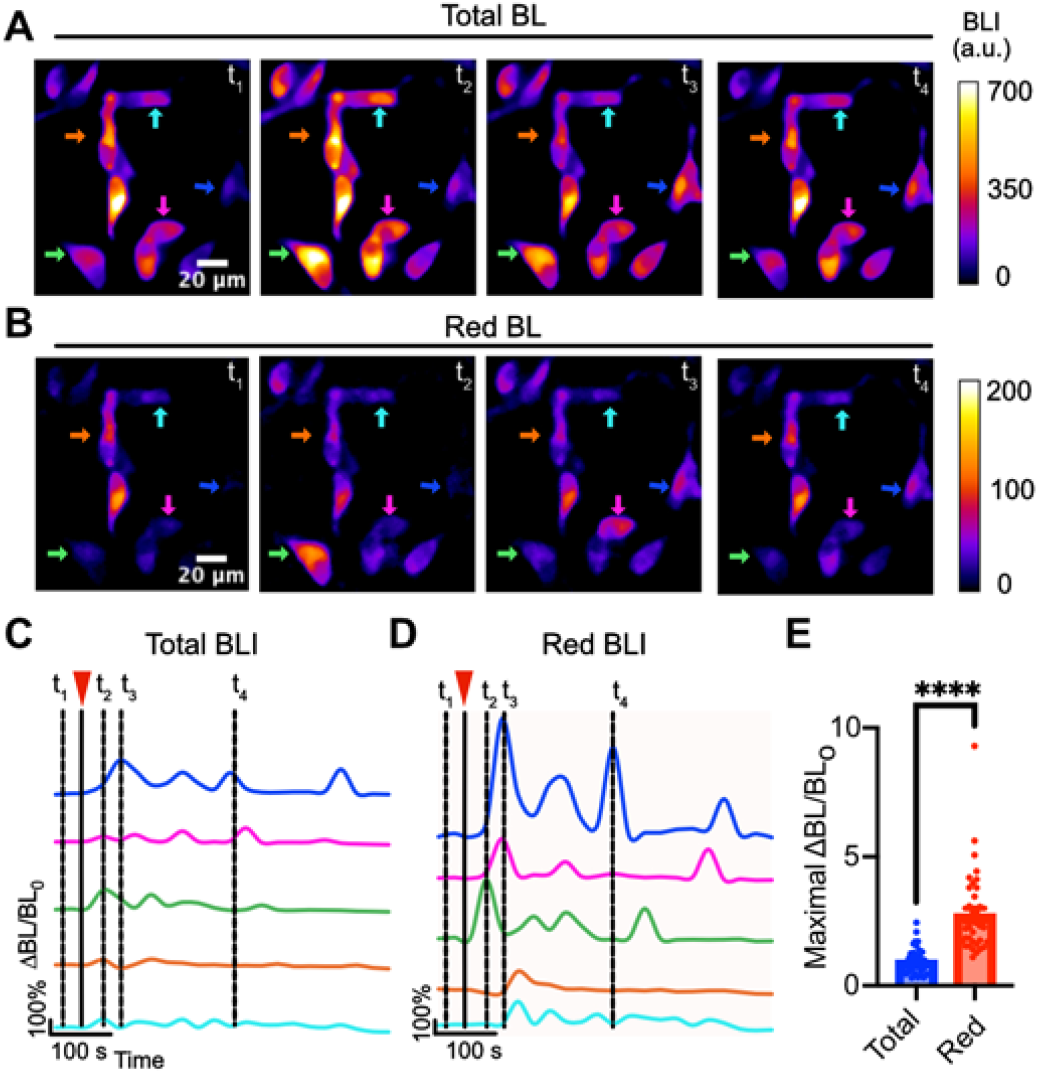
HyBRIC2 resolves histamine-induced Ca^2+^ oscillations in HeLa cells with enhanced red-window contrast. (**A**) Representative pseudocolored images of total bioluminescence intensity from HyBRIC2 expressing HeLa cells at four time points during histamine-induced Ca^2+^ oscillations. (**B**) Corresponding red-window BLI images from the same cells and time points. Colored arrows identify representative cells analyzed in panels C and D. Scale bars, 20 μm. (**C**,**D**) Time-resolved, baseline-normalized changes in total BLI (C) and red-window BLI (D) from the indicated cells. 100% scale bars denote a ΔBL/BL□ value of 1. The red triangles indicate the addition of 20 μM histamine, and the dashed lines mark the time points (*t*_1_-*t*_4_) shown in panels A and B. (**E**) Comparison of the maximal fractional responses (ΔBL/BL□) measured from total and red-window bioluminescence. Bars represent mean ± SEM, and each point represents one cell (*n* = 30 cells). Statistical significance was determined using paired two-tailed *t* tests; ****P < 0.0001.

## DISCUSSION

This study demonstrates that Ca^2+^-dependent luciferase activation and BRET modulation can be coupled within a single genetically encoded indicator to enhance bioluminescence in the red optical window. The HyBRIC architecture combines CaM-M13 regulated LumiLuc activation with TnC-dependent modulation of BRET to mScarlet-I. Ca^2+^ binding therefore increases photon production while also redirecting a greater fraction of the emitted light toward the red acceptor. By allowing these two processes to reinforce each other, the hybrid architecture provides a mechanism for amplifying Ca^2+^-dependent emission above 600 nm. This approach complements recent efforts to improve the brightness, dynamic range, and long-wavelength emission of bioluminescent Ca^2+^-indicators through luciferase and sensor engineering.

The Ca^2+^ titration results are consistent with stepwise activation of the two sensing modules. In HyBRIC, the donor-associated response at 450 nm had an apparent *K*_d_ of approximately 21 nM, whereas the acceptor-associated response at 595 nm had an apparent *K*_d_ of approximately 490 nM. Thus, the higher-affinity CaM-M13 module likely promotes split-luciferase reconstitution and increases photon production at lower Ca^2+^ concentrations, whereas the TnC-dependent transition enhances BRET at higher Ca^2+^ concentrations. This sequence first increases the photon pool available for detection and then redistributes a greater fraction of those photons into the red optical window.

Our study also illustrates that a large *in vitro* response does not necessarily translate into optimal performance in living cells. Although HyBRIC exhibited strong Ca^2+^-dependent changes *in vitro*, its response was substantially smaller in mammalian cells. Our results suggested that the complex multidomain construct may have inefficient protein folding. In addition, the donor-associated apparent *K*_d_ of approximately 21 nM likely caused partial activation of the CaM-M13 regulated catalytic module at resting cytosolic Ca^2+^ concentrations, thereby reducing the fraction of sensors available to respond to stimulation.

Directed evolution using screening conditions that better represented the physiological Ca^2+^ range yielded HyBRIC2. The use of 65 nM and 1.35 μM free Ca^2+^ favored variants with improved response separation over a more relevant concentration range. HyBRIC2 showed a shift in the donor-associated apparent *K*_d_ from approximately 21 to 64 nM while largely retaining the BRET-associated response. This change likely reduced basal catalytic activation and preserved a larger responsive sensor population. Consistent with this interpretation, HyBRIC2 exhibited markedly improved live-cell performance. These findings emphasize the importance of tuning sensor affinity and screening conditions to the intended cellular environment rather than optimizing only the response between Ca^2+^-free and saturating conditions.

From HyBRIC to HyBRIC2, nine mutations were obtained. Although it is difficult to interpret the roles of each mutation in HyBRIC2, the mutations G586E and N594D in the CaM-M13 fragment are located either at the flexible loop between CaM and M13 or at the CaM-M13 interface (**Fig. S4**). We speculate that they may be important for the observed affinity tuning. In addition, other mutations, in particular those in TnC and D553E mutation at the surface of CaM may reduce the crosstalk of the two Ca^2+^ binding domains, leading to the increase robustness of HyBRIC2. As of now, those are mainly speculations, further research is needed to pinpoint the mechanistic root.

Substrate selection was also critical for maximizing red-window contrast. DTZ and 8pyDTZ produced more red-shifted donor emission and greater spectral overlap with mScarlet-I than pyCTZ. However, their red-shifted donor emission also increased bleedthrough into the red window. In contrast, pyCTZ produced lower basal emission above 600 nm and therefore provided the greatest Ca^2+^-dependent contrast, despite its less red-shifted donor spectrum. These results show that maximizing spectral overlap or BRET efficiency alone does not necessarily maximize sensor performance.

HyBRIC2 robustly resolved ACh- and histamine-induced Ca^2+^ dynamics in HEK293T and HeLa cells. The consistently greater contrast in the red-window signal than in total bioluminescence supports the functional benefit of coupling catalytic activation to BRET-mediated spectral redistribution. In fact, the response of HyBRIC2 in the red window to histamine stimulation in HeLa cells was on par with the best available bioluminescent Ca^2+^ indicator as of today (**Table S1**). Nevertheless, the present study was limited to cultured cells, and the suitability of HyBRIC2 for tissue and *in vivo* imaging remains to be established. Its photon output and temporal resolution could also be improved, particularly for monitoring rapid events such as neuronal Ca^2+^ transients. Its high-affinity Ca^2+^-binding CaM-M13 module may need further tunning to better match physiological change. Future engineering to improve folding, maturation, brightness, affinity, kinetics, and substrate delivery may expand the range of biological applications.

In conclusion, we developed HyBRIC and HyBRIC2 by integrating Ca^2+^-dependent luciferase activation and BRET modulation within a single molecular architecture. Pairing the hybrid scaffold with pyCTZ minimized donor bleedthrough and enhanced Ca^2+^-dependent contrast in the red optical window. The two sensors showed large 55-fold Ca^2+^-dependent bioluminescence increase *in vitro*. The further engineered HyBRIC2 performed well in live cells, producing an nearly two-fold red-window response (ΔBL/BL_0_) in acetylcholine-stimulated HEK293T cells and four-fold response in resolving histamine-induced Ca^2+^ oscillations in HeLa cells. These results establish coupling luciferase activation and BRET as a modular strategy for developing high-contrast, long-wavelength bioluminescence biosensors. This modular strategy may be applicable not only to Ca^2+^ indicators but also to bioluminescent biosensors for other cellular analytes.

## AUTHOR INFORMATION

### Author Contributions

H.A. conceived, supervised, and obtained funding for this study. Y.X. engineered and characterized the initial HyBRIC sensor. X.T. performed the directed evolution that yielded HyBRIC2. R.S. recharacterized HyBRIC, further characterized HyBRIC2, and compared the performance of both sensors in mammalian cells. R.S. and Y.X. analyzed data and prepared the initial figures. P.P.S. contributed to the preparation of some luciferins used in this study. R.S., Y.X., and H.A. wrote the manuscript. All authors reviewed and approved the final version of the manuscript.

### Note

University of Virginia filed patent applications covering LumiLuc and sensor configurations described in this manuscript. H.A., Y.X. and X.T. are listed as co-inventors.

## ACKNOWLEDGMENT

The research presented in this publication was supported by the University of Virginia Start-up Fund and the National Institutes of Health grant R01EB035430, R01EB033172, and R01AG077773. ChatGPT was used to paraphrase sentences and rectify grammatical errors.

## For TOC only

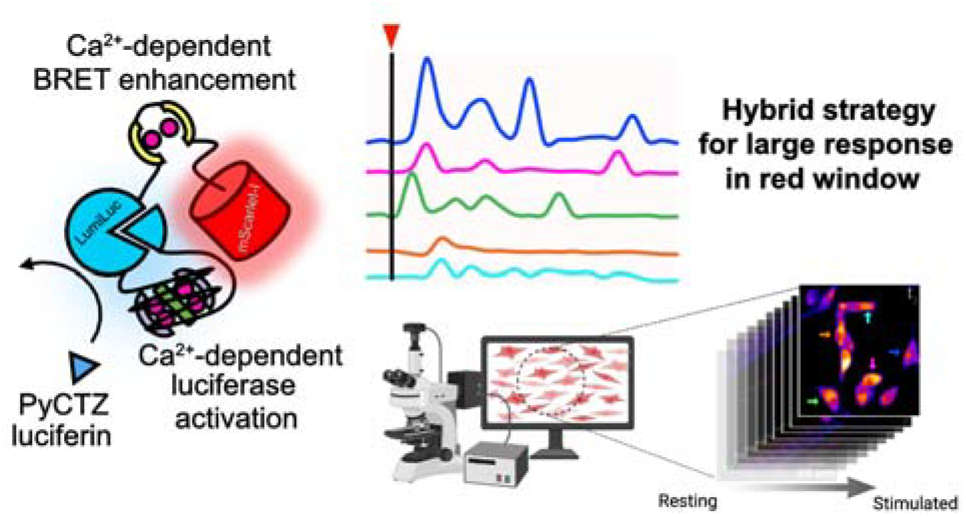

