## Supporting Information for "Coupling Luciferase Activation and BRET Enables High-Contrast Red-Window Calcium Imaging"

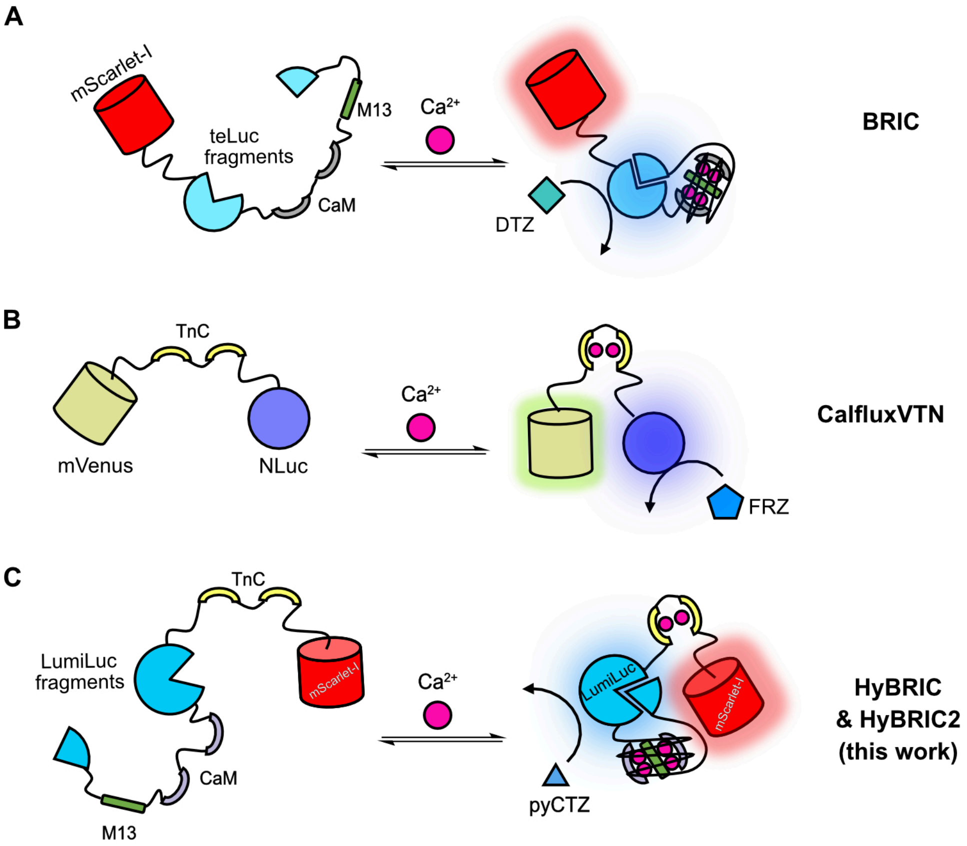


**Fig. S1. Comparison of the $Ca^{2+}$-sensing mechanisms of HyBRIC/HyBRIC2 with those of BRIC and CalfluxVTN.** (**A**) Schematic illustration of the BRIC sensing mechanism. $Ca^{2+}$ binding to the CaM-M13 module promotes reconstitution of the teLuc fragments, thereby increasing luciferase activity and photon output. DTZ provides strong spectral overlap between teLuc emission and mScarlet-I absorption, resulting in efficient BRET; however, $Ca^{2+}$ binding does not directly modulate the BRET process. (**B**) Schematic illustration of the CalfluxVTN sensing mechanism. $Ca^{2+}$ binding to the troponin C (TnC) domain changes the relative geometry of the NLuc donor and mVenus acceptor, thereby modulating BRET efficiency and producing a ratiometric emission response. (**C**) Schematic illustration of the HyBRIC and HyBRIC2 sensing mechanisms. $Ca^{2+}$ binding promotes both reconstitution and activation of the split LumiLuc fragments and TnC-mediated enhancement of BRET to mScarlet-I. These two $Ca^{2+}$-dependent processes collectively increase the sensor response in the red optical window.

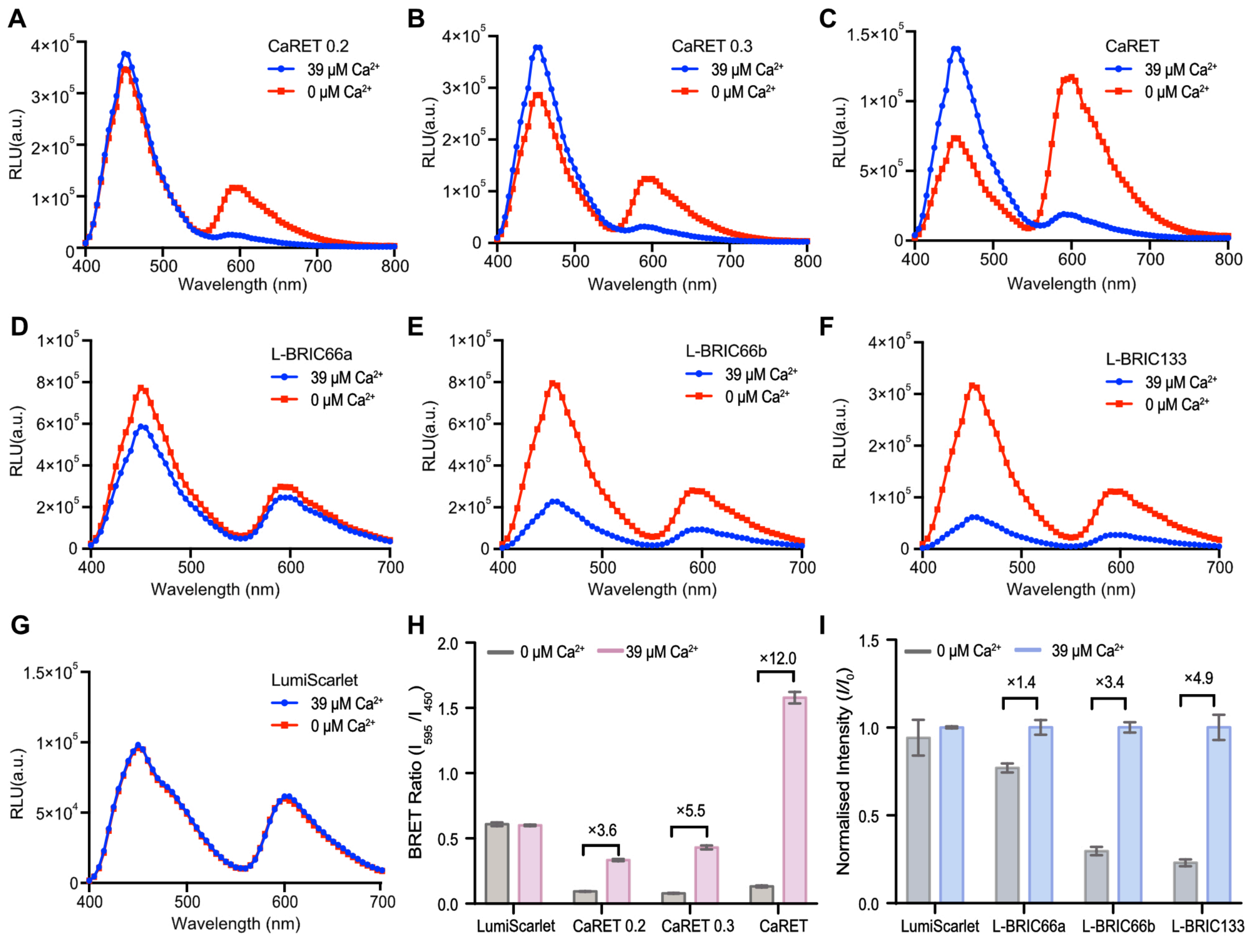


**Fig. S2. Bioluminescence of the ratiometric BRET and luciferase-insertion catalytic modules during HyBRIC engineering.** (**A-C**) Bioluminescence emission spectra of CaRET0.2, CaRET0.3, and the optimized CaRET variant, respectively, showing the progressive enhancement of $Ca^{2+}$-dependent BRET modulation. (**D-F**) Bioluminescence emission spectra of the luciferase-insertion variants L-BRIC66a, L-BRIC66b, and L-BRIC133, respectively, under $Ca^{2+}$-free and $Ca^{2+}$-saturated conditions. (**G**) Bioluminescence emission spectra of the $Ca^{2+}$-insensitive LumiScarlet control. (**H**) Quantification of the BRET ratio ($I_{595}/I_{450}$; ratios of the emission intensity at 595 nm to that at 450 nm) for LumiScarlet and the CaRET variants under $Ca^{2+}$-free and $Ca^{2+}$-saturated conditions. The corresponding $Ca^{2+}$-dependent fold changes are indicated above the brackets. (I) Quantification of the normalized bioluminescence intensity of LumiScarlet and the L-BRIC variants at 450 nm under $Ca^{2+}$-free and $Ca^{2+}$-saturated conditions. The $Ca^{2+}$-dependent fold changes are indicated above the brackets. All measurements were performed using cell lysates under $Ca^{2+}$-free conditions (0 µM $Ca^{2+}$) or $Ca^{2+}$-saturated conditions (39 µM $Ca^{2+}$). In panels H and I, data are presented as mean ± SD from $n$ = 3 technical replicates.

```
         1                                                50
HyBRIC   M V S K G E A V I K E F M R F K V H M E G S M N G H E F E I E G E G E G R P Y E G T Q T A K L K V T
HyBRIC2  M V S K G E A V I K E F M R F K V H M E G S M N G H E F E I E G E G E G R P Y E G T Q T A K L K V T

         51                                               100
HyBRIC   K G G P L P F S W D I L S P Q F M Y G S R A F I K H P A D I P D Y Y K Q S F P E G F K W E R V M N F
HyBRIC2  K G G P L P F S W D I L S P Q F M Y G S R A F I K H P A D I P D Y Y K Q S F P E G F K W E R V M N F

         101                                              150
HyBRIC   E D G G A V T V T Q D T S L E D G T L I Y K V K L R G T N F P P D G P V M Q K K T M G W E A S T E R
HyBRIC2  E D G G A V T V T Q D T S L E D G T L I Y K V K L R G T N F P P D G P V M Q K K T M G W E A S T E R

         151                                              200
HyBRIC   L Y P E D G V L K G D I K M A L R L K D G G R Y L A D F K T T Y K A K K P V Q M P G A Y N V D R R L
HyBRIC2  L Y P E D G V L K G D I K M A L R L K D G G R Y L A D F K T T Y K A K K R V Q M P G A Y N V D R R L

         201                                              250
HyBRIC   D I T S H N E D Y T V V E Q Y E R S E G R K R Q D A S E E E L S E C F R V F D K D G N G F I D R E E
HyBRIC2  D I T F H N E D Y T V V E Q Y E R S E G R K R Q D A S E E E L S E C F R V F D K D G D G F I D R E E

         251                                              300
HyBRIC   F G D I I R L T G E Q L T D E D P D E I F G D S D T D K D G R I D F D E F L K M V E N V Q Y V N I L
HyBRIC2  F G D I I R L T G E Q L S D E D P D E I F G D T D T D K D G R I D F D E F L K M V E N V Q Y V N I L

         301                                              350
HyBRIC   K V F T L G D F V G D W R Q T A G Y N Q A Q V L E Q G G L T S L F Q N L G V S V T P I Q R I V L S G
HyBRIC2  K V F T L G D F V G D W R Q T A G Y N Q A Q V L E Q G G L T S L F Q N L G V S V T P I Q R I V L S G

         351                                              400
HyBRIC   E N G L K I D I H V I I P Y E G L S C D Q M A Q I E K I F K V V Y P V D D H H F K A I L H Y G T L V
HyBRIC2  E N G L K I D I H V N I P Y E G L S C D Q M A Q I E K I F K V V Y P V D D H H F K A I L H Y G T L V

         401                                              450
HyBRIC   I D G V T P N M I D Y F G Q P Y E G I A K F D G K K I T V T G T L I M H D Q L T E E Q I A E F K E A
HyBRIC2  I D G V T P N M I D Y F G Q P Y E G I A K F D G K K I T V T G T L I M H D Q L T E E Q I A E F K E A

         451                                              500
HyBRIC   F S L F D K D G D G T I T T K E L G T V M R S L G Q N P T E A E L Q D M I N E V D A D G N G T I Y F
HyBRIC2  F S L F D K D G D G T I T T K E L G T V M R S L G Q N P T E A E L Q D M I N E V D A D G N G T I Y F

         501                                              550
HyBRIC   P E F L T M M A R K M K D T D S E E E I R E A F R V F D K D G N G Y I S A A Q L R H V M T N L G E K
HyBRIC2  P E F L T M M A R K M K D T D S E E E I R E A F R V F D K D G N G Y I S A A Q L R H V M T N L G E K

         551                                              600
HyBRIC   L T D E E V D E M I R E A D I D G D G Q V N Y E E F V Q M M T A K G G G S K R R W K K N F I A V S A
HyBRIC2  L T E E E V D E M I R E A D I D G D G Q V N Y E E F V Q M M T A K G G E S K R R W K K D F I A V S A

         601                                              650
HyBRIC   A N R F K K I S S S G A L E L W N G N T I I D E R L I N P D G S L L F R V T I N G V T G W R L H E R
HyBRIC2  A N R F K K I S S S G A L E L W N G N T I I D E R L I N P D G S L L F R V T I N G V T G W R L H E R

         651 653
HyBRIC   I L A -
HyBRIC2  I L A -
```

**Fig. S3. Amino acid sequence alignment of HyBRIC and HyBRIC2.** Residues derived from mScarlet-I, TnC, LumiLuc, CaM, and M13 are shown in red, chartreuse, cyan, dark magenta, and orange, respectively. Linker residues are shown in black. Amino acid substitutions in HyBRIC2 relative to HyBRIC are shaded in green.

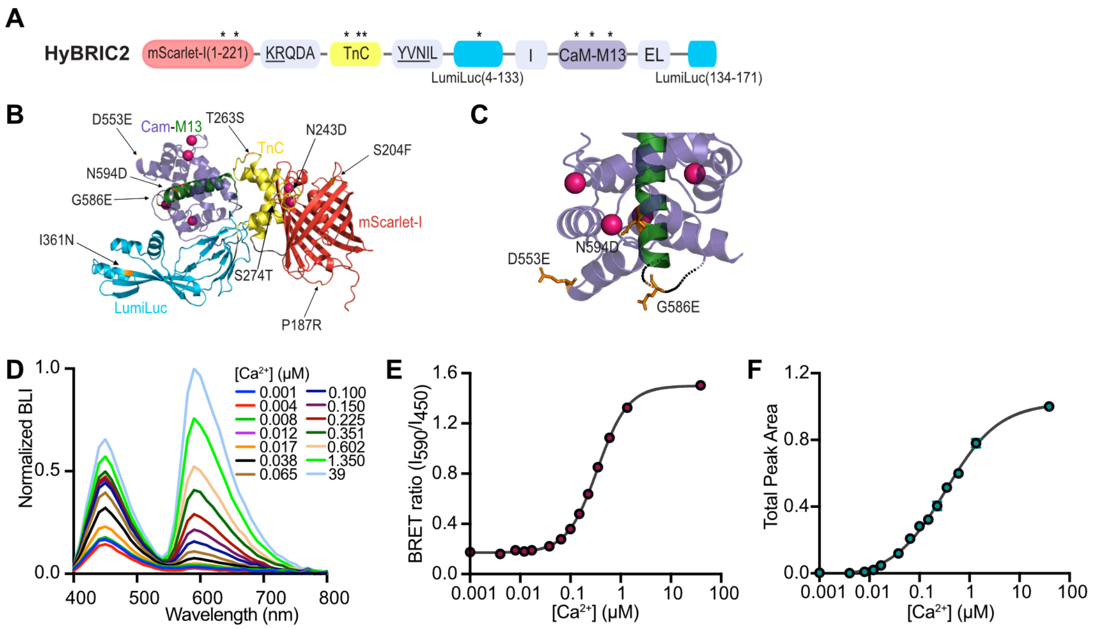


**Figure S4**. **Domain organization, predicted structure, and additional *in vitro* characterization of HyBRIC2.** (**A**) Domain organization of HyBRIC2. Amino acid substitutions relative to HyBRIC are indicated by asterisks above the corresponding domains. (**B**) AlphaFold 3-predicted structure of HyBRIC2 showing the locations of the nine substitutions identified during directed evolution: P187R, S204F, N243D, T263S, S274T, I361N, D553E, G586E, and N594D. (**C**) Enlarged view of the CaM-M13 region. $Ca^{2+}$ ions are shown as magenta spheres, and the D553E, G586E, and N594D substitutions are highlighted as orange sticks. (**D**) Normalized bioluminescence emission spectra of HyBRIC2 over a range of free $Ca^{2+}$ concentrations from 0.001 µM to 39 µM. (**E**) $Ca^{2+}$ titration curve based on the BRET ratio ($I_{595}/I_{450}$). Four-parameter Hill fitting yielded an apparent $K_d$ of 345 nM and a Hill coefficient of 1.4. (**F**) $Ca^{2+}$ titration curve based on total bioluminescence integrated from 400 to 800 nm. Four-parameter Hill fitting yielded an apparent $K_d$ of 335 nM and a Hill coefficient of 0.8. For panels E and F, data are presented as mean ± SEM from $n$ = 3 replicates.

**Table S1.** Characteristics of NanoLuc-derived bioluminescent $Ca^{2+}$ indicators with notable >600 nm red-window emission.

| | **Peak emission (nm)** | ***K*$_d$ (nM)** | **$\Delta$BL/BL$_0$ *in vitro*** | **$\Delta$BL/BL$_0$ in cells** | **Reference** |
|---|---|---|---|---|---|
| **Orange CaMBIs** | 586 | 110-300 | 6 | 0.5$^{\dagger}$ | 1 |
| **BRIC** | 595 | 133 | 5.5 | 0.8$^{\dagger}$ | 2 |
| **eBRIC** | 595 | 2300 | 16 | 3.7$^{\dagger}$ | 3 |
| **CaMBI3-350** | 608 | 350 | 24 | 2.7$^{\&}$ | 4 *(preprint)* |
| **HyBRIC** | 595 | 238 (21 and 490)$^{\ddagger}$ | 54$^{\#}$ | not determined | This work |
| **HyBRIC2** | 595 | 335 (64 and 489)$^{\ddagger}$ | 54$^{\#}$ | 4$^{\dagger\#}$ | This work |

$^{\dagger}$Maximal changes induced with 20 μM histamine in HeLa cells.

$^{\&}$The preprint only reported data for 10 μM histamine in HeLa cells.

$^{\ddagger}$ In parentheses are apparent $K_d$ values derived from 450 and 595 nm emission, recpectively.

$^{\#}$ Dynamic ranges in the red window.
